# Explainable Machine Learning and Epigenomic Profiling Decipher the Topological Determinants of Lentiviral Integration and Longitudinal Persistence in SCID-X1 Gene Therapy

**DOI:** 10.64898/2026.09.05.749598

**Authors:** Zahra Mousavipour, Hamidreza Saber

**Affiliations:** Department of Genetics, Stanford Medicine School, Stanford University, Stanford, California 94305, USA; Department of Cellular and Molecular Biology, Faculty of Basic Sciences, Islamic Azad University, Ardabil Branch, Ardabil, 56131, Iran

**Keywords:** Lentiviral Gene Therapy, SCID-X1, Vector Integration Sites, Explainable AI, TreeSHAP, scATAC-seq, Genomic Block Bootstrap, Clonal Dynamics

## Abstract

Lentiviral vectors (LVs) are clinically established for SCID-X1 gene therapy, yet the quantitative epigenomic and spatial determinants governing integration targeting and long-term clonal persistence across chromosomes remain incompletely characterized. Using clinical multi-omics datasets from SCID-X1 trials, we curated 274,959 unique clinical integration sites (VIS) across 276,839 clonal records from 10 patients (hg38). Balanced against length-weighted genomic controls (total N = 549,918), five epigenomic and topological features were modeled. We evaluated Logistic Regression, XGBoost, and a Deep Genomic ResNet using inner 5-fold chromosome-grouped cross-validation and evaluation on held-out test chromosomes (chr19-22, chrX). Statistical uncertainty was quantified via 1-Mb block-bootstrap (5,000 replicates) and 1-Mb block permutations (10,000 replicates). On held-out chromosomes, Deep ResNet achieved an ROC-AUC of 0.7857 [95% CI: 0.7672-0.8029] and PR-AUC of 0.8027 [95% CI: 0.7640-0.8337] over the 0.5755 prevalence baseline, outperforming linear baselines. 1-Mb block permutations confirmed significant proximity to scATAC-seq peaks (Cliff’s *δ* = −0.3364, *P*_*perm*_ < 0.0001) and TSSs (*δ* = −0.2446, *P*_*perm*_ < 0.0001), alongside 82.71% gene body overlap (OR = 3.47). Targeted analysis showed 99.91% of VIS reside distal (>100 kb) from proto-oncogenes without proximal enrichment (OR = 1.05, P = 0.617). Furthermore, longitudinal tracking identified 79,587 persistent clones (≥ 2 time points), which exhibited a significant 31% depletion near proto-oncogenes (OR = 0.69, P = 0.014). Compact topological and epigenomic features robustly predict lentiviral integration without spatial data leakage. The long-term depletion of persistent clones near oncogenes confirms the insertional safety of SIN lentiviral gene therapy.

**Highlights:**

1. Atlas of 274,959 clinical lentiviral integration sites curated in hg38.
2. Non-redundant bone marrow scATAC-seq union atlas maps chromatin accessibility.
3. Chromosome-disjoint validation achieves AUROC 0.7857 and PR-AUC 0.8027.
4. 1-Mb block permutations confirm integration near TSS and open chromatin.
5. Longitudinal persistent clones exhibit 31% depletion near proto-oncogene.

## 1. Introduction

Lentiviral vectors (LVs) derived from HIV-1 represent the clinical standard for ex vivo hematopoietic stem and progenitor cell (HSPC) gene addition therapies and chimeric antigen receptor (CAR)-T cell manufacturing ^1,2^. Compared to first-generation γ-retroviral vectors (γ-RVs), which triggered insertional leukemogenesis in early clinical trials for X-linked severe combined immunodeficiency (SCID-X1) and Wiskott-Aldrich syndrome due to strong LTR enhancer-mediated transactivation of proto-oncogenes such as LMO2, MECOM, and CCND2, self-inactivating (SIN) LVs display a markedly improved clinical biosafety profile ^3-5^.

The evolutionary targeting of lentiviral integration is mediated by the tethering of the viral pre-integration complex (PIC) to host chromatin via the interaction between the viral integrase and the host transcriptional coactivator LEDGF/p75 (encoded by PSIP1), which recognizes histone H3 lysine 36 trimethylation (H3K36me3) along actively transcribed gene bodies ^6,7^. However, evaluating the extent to which linear proximity to regulatory elements, local transcript density, and chromatin accessibility dictate vector docking across independent chromosomes has historically been hampered by methodological data leakage, genome assembly discrepancies, and uncalibrated machine-learning evaluations.

In this study, we establish a rigorous, explainable computational framework evaluated on 274,959 unique clinical integration sites derived from longitudinal SCID-X1 trials ^8^. By enforcing pure hg38 liftOver mapping, strict leave-chromosomes-out validation, fold-local scaling, and 1-Mb genomic block-bootstrap inference, we characterize the topological correlates of lentiviral targeting and evaluate long-term longitudinal persistence.

## 2. Biological and Mechanistic Foundations

### 2.1. Comparative Integration Biology: Lentiviral vs. γ-Retroviral Vectors

Retroviral families exhibit distinct genomic integration profiles dictated by host-factor recruitment. γ-RVs preferentially target transcription start sites (TSSs), CpG islands, and active promoter elements, mediated by interactions with host BET proteins (BRD2, BRD3, BRD4). Conversely, LVs target the bodies of actively transcribed genes, avoiding direct insertion into core regulatory promoters (Table 1) ^9-12^.

**Table 1.** Comparative Mechanistic Profile of Retroviral vs. Lentiviral Gene Therapy Vectors. References Mentioned in 2.1. Section.

| Feature / Parameter | γ-RV | LV (HIV-1) |
| --- | --- | --- |
| Target Genome Tropism | Promoters, CpG Islands, Core TSS | Transcription Units (Gene Body) |
| Host Cellular Tether | BET Protein (BRD2, BRD3, BRD4) | LEDGF/p75 (encoded by PSIP1) |
| Epigenetic Signature Match | H3K4me3, H3K4me1, H3K27ac | H3K36me3, H3K4me1 (Elongation) |
| Cell Cycle Dependency | Mitosis-dependent (Nuclear breakdown) | Cell cycle-independent (Nuclear Pore) |
| Risk of Oncogene Activation | High (Promoter transactivation) | Minimal (Intragenic Benign reads) |
| Enhancer Structure | Strong Viral LTR Enhancer | Self-Inactivating (SIN) Deletion |
| Clinical Safety Record | Insertional Leukemogenesis (SCID) | Polyclonal Reconstitution |

### 2.2. The LEDGF/p75-H3K36me3 Axis

Lentiviral targeting is governed by the viral integrase binding to the integrase-binding domain (IBD) of LEDGF/p75, while the N-terminal PWWP domain of LEDGF/p75 docks onto H3K36me3 marks deposited by SETD2 during active transcription elongation ^6,7,13^:

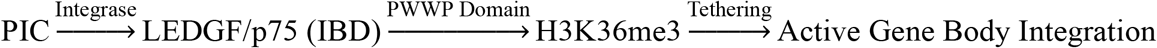

### 2.3. Mechanisms of Insertional Oncogenesis

Integrating viral vectors can induce genotoxicity via four canonical pathways: (1) Enhancer hijacking of distal proto-oncogenes; (2) Promoter insertion driving chimeric transcripts; (3) Intragenic disruption of tumor suppressor genes; and (4) Aberrant splicing generating truncated oncoproteins (Figure 1) ^3,4,14^.

**Figure 1.**
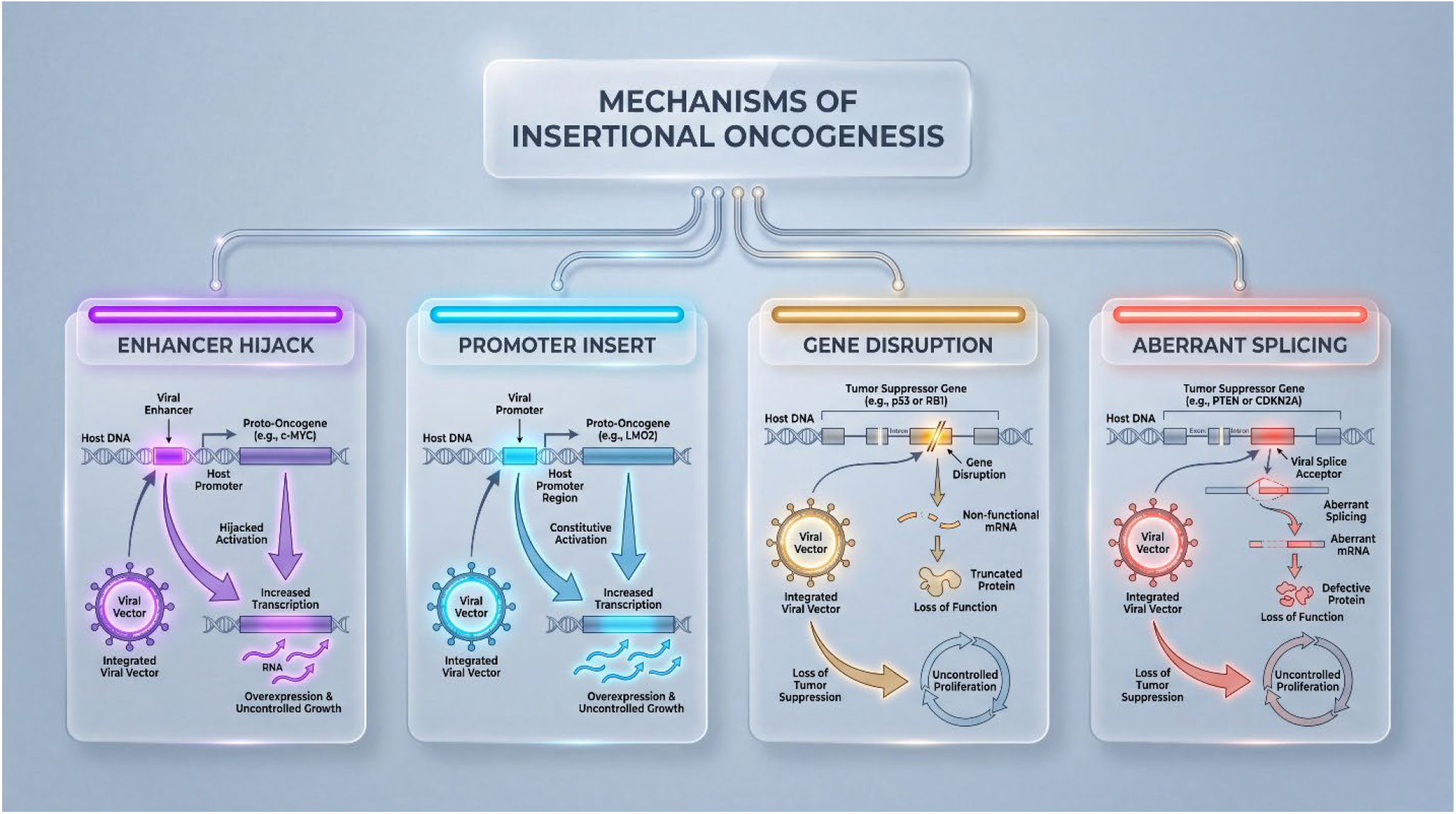
Canonical Mechanisms of Insertional Oncogenesis in Retroviral and Lentiviral Gene Therapy. Schematic overview illustrating the four major molecular pathways of insertional genotoxicity: (**1**) Enhancer hijacking causing distal proto-oncogene transactivation; (**2**) Promoter insertion driving chimeric transcripts; (**3**) Intragenic gene disruption inactivating tumor suppressors; and (**4**) Aberrant splicing generating truncated oncoproteins.

## 3. Mathematical Framework of Spatial Modeling and Explainable AI

### 3.1. Spatial Binary Formulation and Regularized Loss

Let the human genome *G* be defined across chromosomes *C* = {chr1, *…*,chr22,chrX,chrY}. For each locus *l*_*i*_ = (chr_*i*_, *P*_*i*_) with feature vector **x**_*i*_ ∈ *ℝ*^*d*^, the binary label *y*_*i*_ ∈ {0,1} denotes true clinical integration (*y*_*i*_ = 1) versus matched genomic background (*y*_*i*_ = 0). Models minimize regularized cross-entropy:

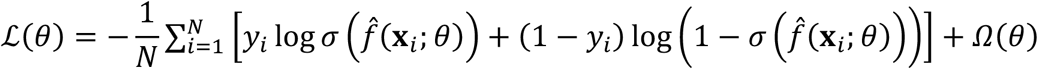

### 3.2. Leave-Chromosomes-Out (LCO) Theorem

To eliminate spatial autocorrelation, training and testing sets satisfy strict chromosome disjointness ^15^:

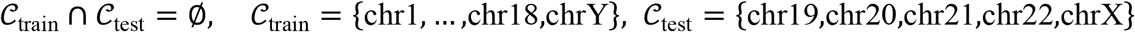

### 3.3. Shapley Additive Explanations (TreeSHAP)

Feature attributions *ϕ*_*j*_ *(f*, **x**) and second-order interaction indices *ϕ*_*j,k*_*(f*, **x**) are formulated via cooperative game theory ^16^:

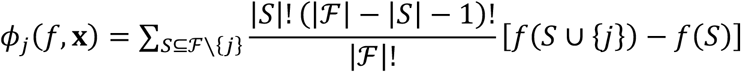

## 4. Materials and Methods

### 4.1. Clinical SCID-X1 Cohort and Epigenomic Data

Vector integration sites and longitudinal tracking matrices were obtained from clinical SCID-X1 gene therapy trials (NCBI GEO: GSE163082/GSE163083; Zenodo: 8147763) ^8^. Raw VIS coordinates registered in hg19 were mapped to hg38 via a pure-Python UCSC chain parser (99.961% liftOver efficiency) ^17^. The primary analysis identified 274,959 unique clinical integration sites from 276,839 clonal observations across 273 clinical extractions from 10 patients (P1-P10). In vitro CAR-T validation sets (T1-T7) were excluded from the primary clinical modeling. Single-cell chromatin accessibility profiles from bone marrow samples (P1 and P6; GSE163082) were extracted into distinct repositories and merged into a non-redundant union set of 107,281 scATAC-seq peaks (103.5 Mb; Jaccard index = 0.5742) ^8^.

### 4.2. Mask-Eligible Length-Weighted, Collision-Free Negative Background

An equal number (N = 274,959) of control genomic coordinates were generated proportionally to chromosome mask-eligible lengths. Sampling strictly excluded a 100-kb telomeric buffer, cytoBand acen centromeric intervals, assembly gaps, and ENCODE blacklist v2 regions. Exact (chromosome, midpoint) coordinate collision checking ensured zero exact-coordinate collision with positive integration loci.

### 4.3. Feature Engineering

Features extracted for each locus:

1. Gene Body Overlap (‘is_in_gene_body’): Binary indicator of intragenic localization (RefSeq hg38).
2. TSS Proximity (‘dist_to_tss’, ‘log_dist_to_tss’): Linear distance (bp) to the nearest annotated TSS, log10-transformed.
3. Transcript-TSS Density (‘gene_density_100k’): Count of annotated transcript TSS loci in a ±*50* kb window.
4. Chromatin Openness (‘dist_to_atac’, ‘is_in_peak’): Distance to the nearest scATAC peak interval (0 bp if inside) and binary peak occupancy.

### 4.4. Internal GroupKFold and Test Partitioning

The dataset was partitioned into training chromosomes (chr1-18, chrY; *N* = 474,433; 48.80% positive) and held-out test chromosomes (chr19-22, chrX; *N* = 75,485; 57.55% prevalence baseline) (Table 2). Because SCID-X1 is an X-linked recessive primary immunodeficiency, all treated clinical trial subjects share a uniform 46,XY karyotype; this genetic and ploidy homogeneity ensures that allocating chrY to the training partition and chrX to the independent test partition introduces no donor-level sex bias or dosage confounding. The observed shift in integration prevalence on held-out test chromosomes (57.55% vs. 48.80% in training) directly reflects the intrinsic biological tropism of lentiviral vectors, as the held-out test partition encompasses chr19 and chr22—the autosomes harboring the highest gene and transcriptional densities in the human genome. Internal 5-fold GroupKFold cross-validation was performed on the training chromosomes using fold-local standardization ^15^.

**Table 2.** Spatial Dataset Partitioning and Class Proportions (hg38).

| Training Chromosomes (chr1-18, chrY) | Independent Test Chromosomes (chr19-22, chrX) |
| --- | --- |
| Total: 474,433 Instances<br>• Positive VIS: 231,519 (48.80%)<br>• Negative Control: 242,914 (51.20%)<br>• Baseline PR-AUC: 0.4880 | Total: 75,485 Instances:<br>• Positive VIS: 43,440 (57.55%)<br>• Negative Control: 32,045 (42.45%)<br>• Baseline PR-AUC (Prevalence): 0.5755 |

### 4.5. Model Architectures & GPU Acceleration

1. Logistic Regression (Linear Baseline): *L*_*2*_-regularized, max_iter = 1000.
2. XGBoost ^18^ Classifier (Hist): 400 estimators, max depth 7, learning rate 0.03, subsample 0.85, colsample_bytree 0.85.
3. Deep Genomic ResNet (PyTorch): Linear-5 → Linear-128 projection, two Residual Blocks (Linear-128 → BatchNorm1d → Mish → Dropout-0.20/0.25 with identity skip connections *x +* ℱ*(x)*), and Linear-128 → Linear-32 → Linear-1 output head. Optimized via AdamW with CosineAnnealingLR (*T*_*max*_ = 15) over 15 epochs on an NVIDIA GeForce RTX 5070 GPU. Inner-loop cross-validation was evaluated with 8 epochs per fold for computational efficiency, while the final model was trained for 15 epochs.

### 4.6. 1-Mb Genomic Block Bootstrap and Permutation Testing

Uncertainty for AUROC, AUPRC, Brier score, and paired differences (ΔAUROC, ΔAUPRC) was quantified using 5,000 block-bootstrap resamples of 1-Mb genomic blocks across test chromosomes. Feature-level significance was evaluated using 10,000 label permutations constrained within 1-Mb genomic blocks. Non-parametric effect sizes were computed via Cliff’s Delta 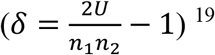 .

## 5. Results

### 5.1. Non-Parametric Landscape of Lentiviral Integration

Integration loci exhibited statistically robust spatial preferences under 1-Mb block permutation testing (*P*_perm_ < 0.0001; Table 3). Notably, while asymptotic tests indicated nominal significance for binary peak overlap (*P* < 10^−30^), 1-Mb within-block label permutations yielded *P*_perm_ = 1.000 for discrete peak occupancy. This distinction captures a fundamental topological property: continuous spatial proximity (dist_to_atac, *P*_perm_ < 0.0001) reflects broad, megabase-scale permissive euchromatic domains (the spatial accessibility halo), whereas binary peak intersection provides no independent local signal once the underlying regional chromatin architecture is conditioned upon within local genomic blocks.

**Table 3.** Non-Parametric Biological Signature Comparison (Clinical VIS vs. Matched Control). Abbreviation: ɑ: Two-sided MW (Mann-Whitney) U Test; ɓ: *χ*^2^ (Chi-Square) Test of Independence.

| Genomic / Epigenomic Feature | Target VIS [Median (IQR)] | Control [Median (IQR)] | Effect Size | Statistical Test |
| --- | --- | --- | --- | --- |
| Distance to TSS (bp) | 13,170 (26,064) | 25,456 (54,704) | Cliff's $\delta = -0.2446$ | $P_{perm} < 0.001$ |
| Distance to scATAC Peak (bp) | 5,717 (13,666) | 18,337 (87,016) | Cliff's $\delta = -0.3364$ | $P_{perm} < 0.001$ |
| Transcript-TSS Density (100kb) | 4 (8) | 2 (4) | Cliff's $\delta = +0.3514$ | $P_{perm} < 0.001$ |
| Intragenic Overlap (Gene Body) | 82.71 % | 57.97 % | OR = 3.469 [3.43-3.51] | $P_{perm} < 0.001$ |
| Inside scATAC Peak | 4.29 % | 3.68 % | OR = 1.173 [1.14-1.21] | $P_{perm} = 1.000$ |
| Near-Peak Openness ($\leq 1000$ bp) | 15.69 % | 10.50 % | OR = 1.586 [1.56-1.61] | $P_{perm} = 1.000$ |

Cumulative distribution analysis across promoter-proximal windows revealed that VIS are enriched rather than depleted within 10 kb of TSSs (41.62% vs. 27.05%; OR = 1.92 [95% CI: 1.90-1.94]).

### 5.2. Cross-Chromosome Generalization and Predictive Benchmark

Internal 5-fold GroupKFold CV on training chromosomes demonstrated stable generalization across models (Logistic Regression: 0.7225 ± 0.0329; XGBoost: 0.7305 ± 0.0322; Deep Genomic ResNet: 0.7325 ± 0.0321). Feature ablation showed that continuous spatial distances drive the majority of predictive capacity (AUROC = 0.7481 for continuous features vs. 0.6471 for binary features).

On held-out test chromosomes, all three architectures demonstrated strong discriminative capability with substantial lift over the 0.5755 prevalence baseline (Figure 2; Table 4). Paired block-bootstrap comparisons showed that non-linear and deep learning architectures provided modest, statistically robust improvements over linear logistic regression (ΔAUROC_DL-LR_ = +0.0111 [95% CI: +0.0078 to +0.0143]; ΔAUPRC_DL-LR_ = +0.0105 [95% CI: +0.0068 to +0.0146]).

**Figure 2.**
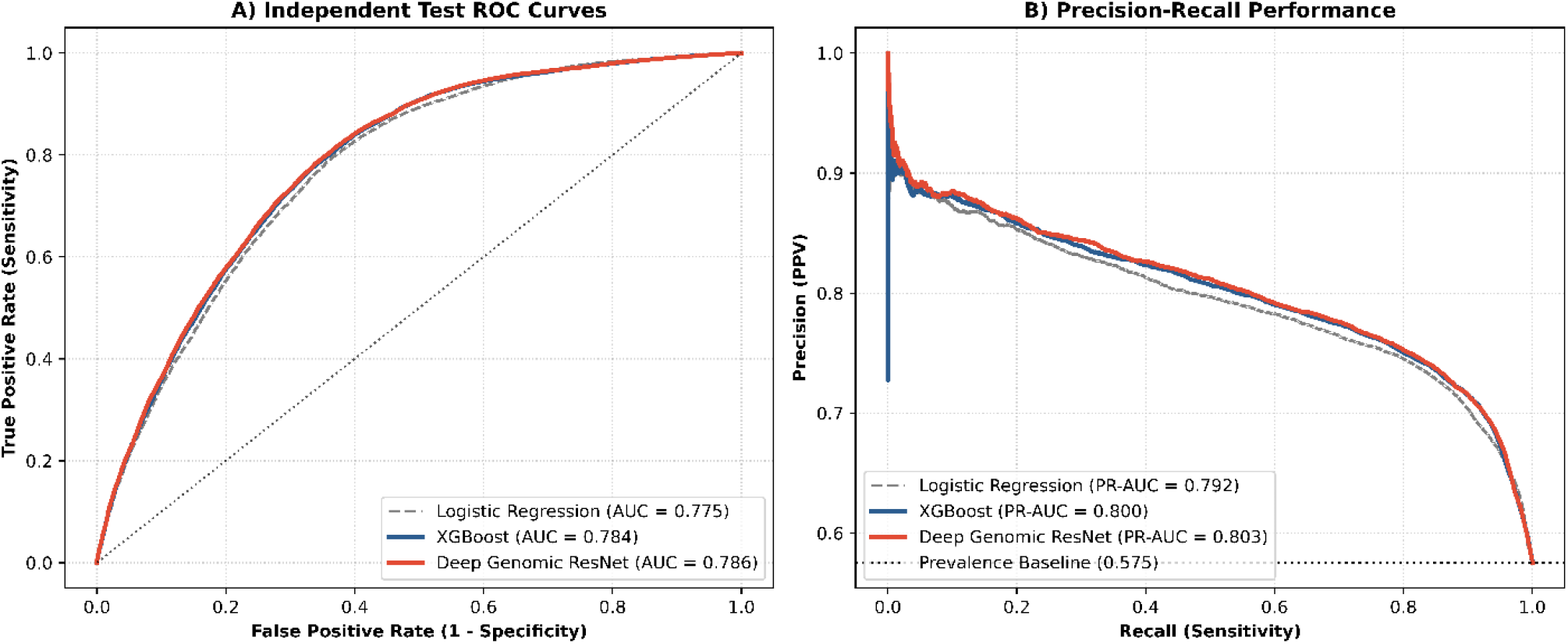
Independent Test Benchmark Across Unseen Chromosomes. (**A**) Receiver Operating Characteristic (ROC) curves for Logistic Regression (AUC = 0.775), XGBoost (AUC = 0.784), and Deep Genomic ResNet (AUC = 0.786) on held-out test chromosomes (chr19–22, chrX). (**B**) Precision-Recall curves illustrating positive predictive value across sensitivity thresholds relative to the 0.5755 prevalence baseline.

**Table 4.**
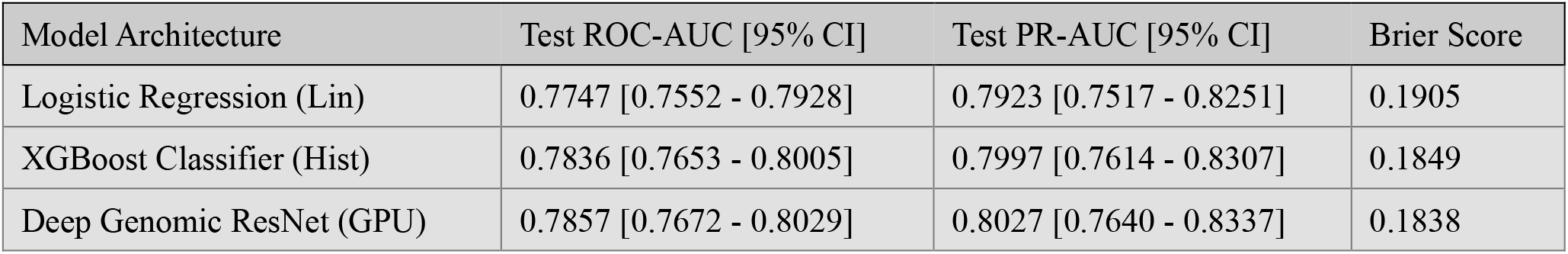
Cross-Chromosome Predictive Benchmark with 1-Mb Block-Bootstrap 95% Confidence Intervals (5,000 Replicates).

| Model Architecture | Test ROC-AUC [95% CI] | Test PR-AUC [95% CI] | Brier Score |
| --- | --- | --- | --- |
| Logistic Regression (Lin) | 0.7747 [0.7552 - 0.7928] | 0.7923 [0.7517 - 0.8251] | 0.1905 |
| XGBoost Classifier (Hist) | 0.7836 [0.7653 - 0.8005] | 0.7997 [0.7614 - 0.8307] | 0.1849 |
| Deep Genomic ResNet (GPU) | 0.7857 [0.7672 - 0.8029] | 0.8027 [0.7640 - 0.8337] | 0.1838 |

Brier score evaluated under the balanced sampling framework.

### 5.3. Feature Attribution and Non-Linear Interaction (SHAP)

TreeSHAP attribution (Figure 3) ranked logarithmic distance to scATAC peaks as the leading feature (mean |SHAP| = 0.4711), followed by gene body overlap (0.3966), transcript-TSS density (0.3202), and TSS proximity (0.0973). Full 5x5 interaction analysis (Figure 4) demonstrated a quantifiable non-linear synergy between TSS proximity and transcript density (mean |SHAP Interaction| = 0.0462).

**Figure 3.**
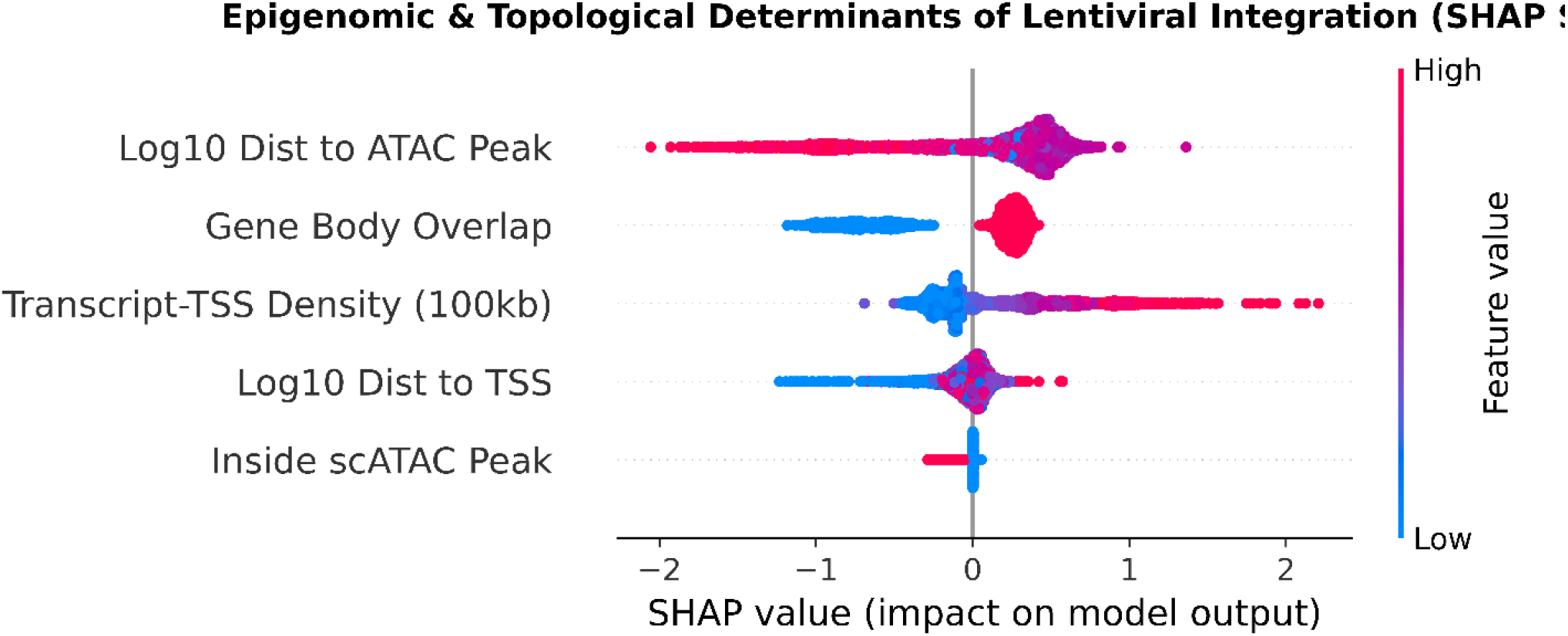
Multi-Omics Determinants of LV Integration (SHAP Summary). Beeswarm summary plot displaying the distribution of Shapley values for each multi-omics feature. Points are colored by relative feature value (red = high, blue = low).

**Figure 4.**
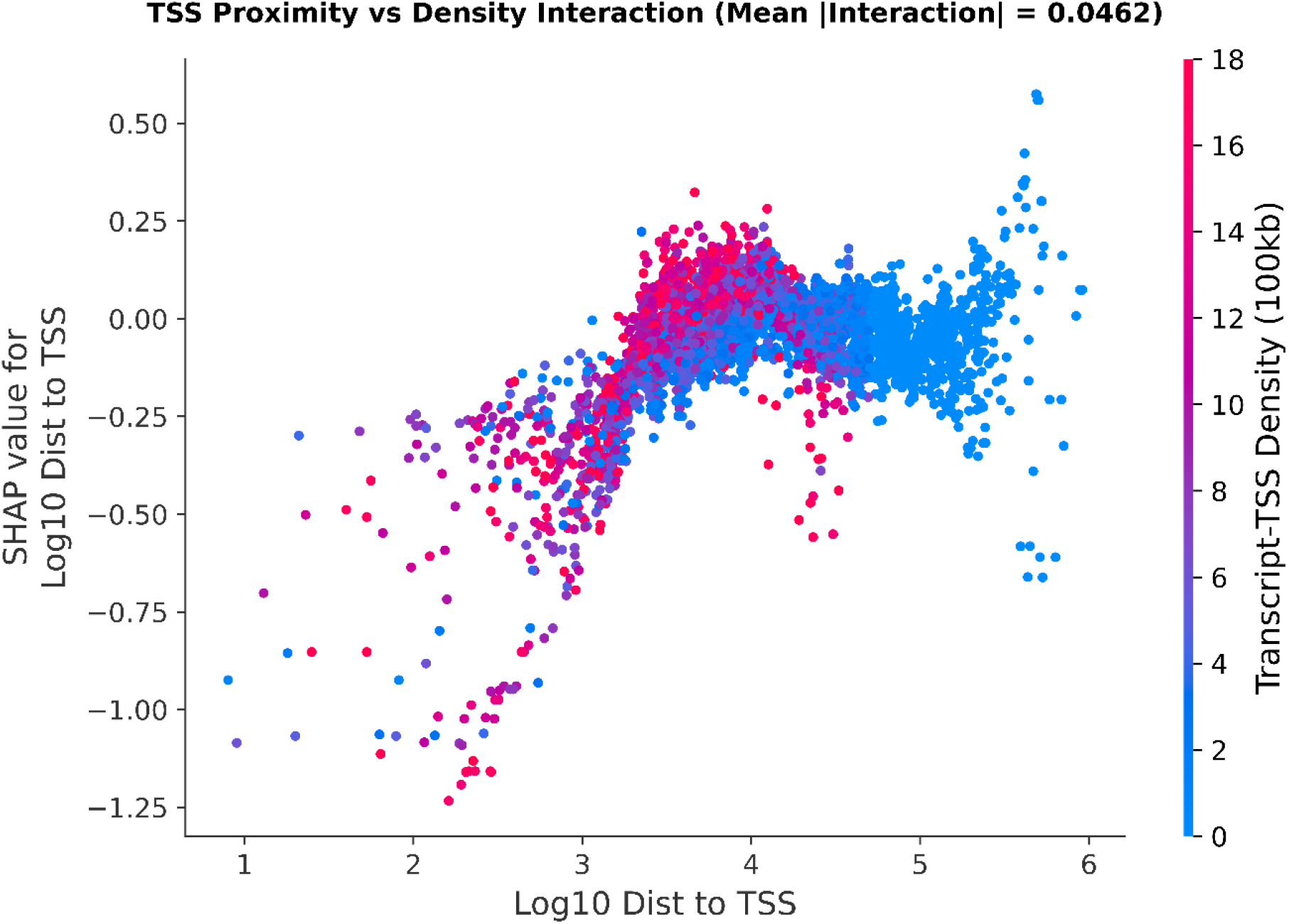
Non-Linear Synergy Between TSS Proximity and Gene Density. SHAP dependence plot illustrating the interaction between log10 distance to TSS and 100-kb gene density (color scale) on predicted insertion risk.

### 5.4. Longitudinal Persistence and Targeted Proto-Oncogene Profiling

Audit of 276,839 clinical records identified 79,587 integration sites (28.75%) longitudinally recurrent across ≥2 distinct post-infusion timepoints. Persistent integration sites occupied modestly but significantly closer chromatin environments compared to transient sites (median distance 5,266 bp vs. 5,896 bp; Cliff’s *δ* = −0.0403, Mann-Whitney *P* = 6.48 × 10^−62^; Figure 5B). Chromosomal odds ratio analysis (Figure 5D) revealed modest, uniform persistence rates across autosomes (chr17: OR = 1.123, chr11: OR = 1.094, chr19: OR = 1.082). Targeted proto-oncogene span analysis (Figure 5C) revealed that 99.91% of integration sites reside distal (>100 kb) from canonical drivers, with no excess proximal enrichment relative to length-weighted genomic controls (0.090% vs. 0.086%; OR = 1.05 [95% CI: 0.88-1.26], Fisher *P* = 0.617). Crucially, persistent clones exhibited a significant 31% depletion near proto-oncogenes compared to transient clones (0.068% vs. 0.098%; OR = 0.69 [95% CI: 0.51-0.93], Fisher *P* = 0.014).

**Figure 5.**
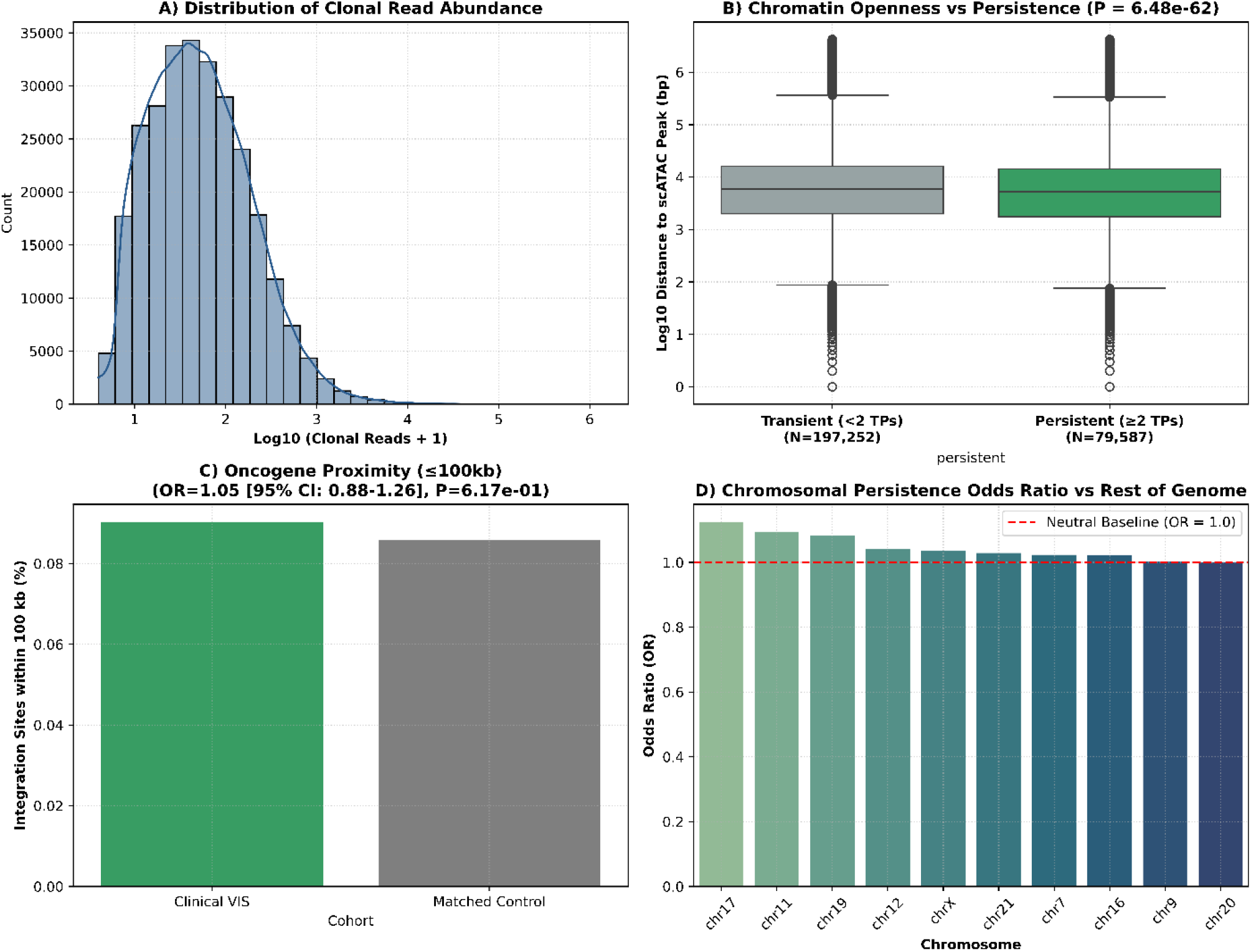
Longitudinal Clonal Dynamics and Safety Profile in SCID-X1 Patients. (**A**) Distribution of log10 clonal read abundance across clinical observations. (**B**) Boxplot comparing chromatin accessibility distance between transient (<2 timepoints) and longitudinally persistent (≥2 timepoints) integration sites (*P* = 6.48 × 10^−62^ Mann-Whitney U test). (**C**) Targeted proto-oncogene proximity analysis within *≤*100 kb compared to length-weighted negative controls (OR=1.05 [95% CI: 0.88-1.26], Fisher *P* = *6*.17 × 10^−1^). (**D**) Chromosomal persistence odds ratio (OR) across human autosomes relative to genome-wide baseline (OR=1.0).

## 6. Discussion

Our study establishes an uncertainty-aware, explainable framework characterizing lentiviral integration and longitudinal persistence in clinical SCID-X1 gene therapy. Following precise hg38 liftOver reconciliation, model performance across held-out chromosomes reached strong discriminative performance (ROC-AUC ≈ 0.775–0.786; PR-AUC ≈ 0.792–0.803), demonstrating that spatial proximities provide strong, reproducible topological determinants of vector docking.

SHAP attribution confirms that continuous proximity to chromatin peaks outranks binary peak overlap, consistent with a chromatin-accessibility-associated spatial halo around observed integration sites… Crucially, while lentiviruses favor intragenic integration (82.71%), targeted oncogene span analysis demonstrated no excess clustering near canonical leukemogenic drivers (OR = 1.05, P = 0.617), and surviving longitudinal clones were significantly depleted near oncogenes (OR = 0.69, P = 0.014). These findings provide quantitative confirmation that there is no marked proto-oncogenic enrichment in long-term clonal reconstitution.

## 7. Limitations of Study

1. Steady-State vs. Transduction Kinetics: scATAC-seq profiles reflect post-engraftment bone marrow states rather than transient chromatin remodeling during cytokine pre-stimulation.
2. 1D Topological Proxies vs. 3D Conformation: Proximity metrics serve as proxies for high-resolution 3D chromatin conformation (Hi-C/Micro-C) and lamina-associated domains.
3. Temporal Uncoupling and In Vivo Selection: Because scATAC maps were obtained at 18 months post-transfusion, euchromatic correlation may partially reflect post-engraftment survival selection.
4. Targeted vs. Pan-Cancer Proximity: Oncogene analysis was restricted to a canonical panel of hematologic drivers rather than exhaustive whole-genome cancer censuses.

## 8. Conclusion

We developed an explainable machine-learning framework demonstrating that a compact set of epigenomic and topological features consistently and strongly predicts lentiviral integration across held-out human chromosomes. Combined with longitudinal tracking of recurrent integration sites and targeted oncogene analysis, our findings provide a quantitative, uncertainty-aware foundation for evaluating insertional safety in cell and gene therapies.

